# Use of multi-flip angle measurements to account for transmit inhomogeneity and non-Gaussian diffusion in DW-SSFP

**DOI:** 10.1101/861880

**Authors:** Benjamin C. Tendler, Sean Foxley, Moises Hernandez-Fernandez, Michiel Cottaar, Connor Scott, Olaf Ansorge, Karla Miller, Saad Jbabdi

## Abstract

Diffusion-weighted steady-state free precession (DW-SSFP) is an SNR-efficient diffusion imaging method. The improved SNR and resolution available at ultra-high field has motivated its use at 7T. However, these data tend to have severe B_1_ inhomogeneity, leading not only to spatially varying SNR, but also to spatially varying diffusivity estimates, confounding comparisons both between and within datasets. This study proposes the acquisition of DW-SSFP data at two-flip angles in combination with explicit modelling of non-Gaussian diffusion to address B_1_ inhomogeneity at 7T. DW-SSFP datasets were acquired from five fixed whole human post-mortem brains with a pair of flip angles that jointly optimize the diffusion contrast-to-noise across the brain. We compared one and two flip-angle DW-SSFP data using a diffusion tensor model that incorporates the full DW-SSFP Buxton signal model. The two-flip angle data were subsequently fitted using a modified DW-SSFP signal model that incorporates a Gamma distribution of diffusivities. This allowed us to generate tensor maps at a single, SNR-optimal effective b-value yielding more consistent SNR across tissue, in addition to eliminating the B_1_ dependence on diffusion coefficients and orientation maps. Our proposed approach will allow the use of DW-SSFP at 7T to derive diffusivity estimates that have greater interpretability, both within a single dataset and between experiments.

**Highlights:**

- B_1_ inhomogeneity at 7T leads to spatially varying SNR & ADC estimates in DW-SSFP
- 2-flip angle DW-SSFP data can address B_1_ effects in a cohort of post-mortem brains
- Our approach reduces degradations in PDD estimates & improves whole brain coverage
- Our approach provides a means to define ADCs at an SNR-optimal effective b-value

## Introduction

Diffusion imaging of post-mortem human brains has important applications for both validating diffusion contrast mechanisms through comparison with microscopy and achieving very high-resolution data with long scan times. However, post-mortem diffusion imaging presents significant challenges due to changes in tissue properties related to death and fixation. Unfavorable reductions in T_1_, T_2_ and diffusion coefficient have been observed in fixed tissue using a variety of fixation methods (Blamire et al., 1999; D’Arceuil and de Crespigny, 2007; Dawe et al., 2009; Shepherd et al., 2009; Yong-Hing et al., 2005).

One method to overcome these changes is to utilize an imaging strategy that allows for fast acquisition of the MR signal to overcome the losses associated with the shortened T_2_ values. We have previously proposed the use of diffusion-weighted steady-state free precession (DW-SSFP) for post-mortem imaging due to its ability to achieve robust signal and strong diffusion contrast in short-T_2_ species (McNab et al., 2009). The high signal-to-noise (SNR) efficiency of DW-SSFP compared to diffusion-weighted spin echo (DW-SE) acquisitions enables improvements in the quality of both diffusion tractography and estimates of multiple fiber populations at 3T (Miller et al., 2012), motivating its use in post-mortem samples (Vasung et al., 2019; Wilkinson et al., 2016).

Ultra-high field scanners have potential to enable further gains in spatial resolution, with DW-SSFP providing a valuable tool for addressing the even shorter T_2_ values at 7T and above (Foxley et al., 2014a). However, DW-SSFP data acquired at 7T are compromised by B_1_ inhomogeneity (Fig. 1a). This presents us with a challenge: unlike other diffusion imaging sequences, both the signal (Fig. 1b) and diffusion attenuation (Fig. 1c) in DW-SSFP are sensitive to flip angle (Buxton, 1993).

**Figure 1:**
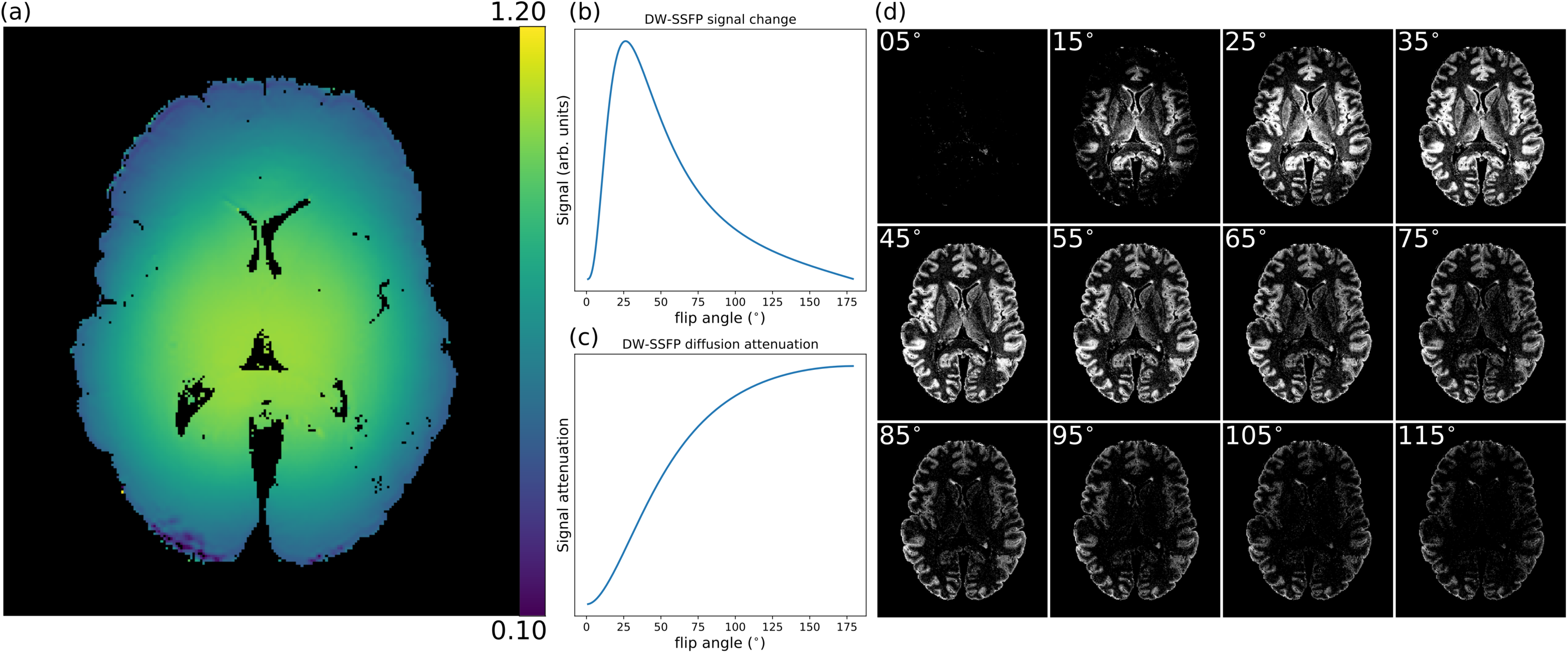
B_1_ inhomogeneities at 7T. (a) A single slice of a B_1_ map estimated using the method described in (Yarnykh, 2007) obtained over a whole post-mortem human brain sample at 7T. B_1_ decreases smoothly as we approach the edge of the brain. The signal (b) and diffusion attenuation (c) have a strong flip angle dependence in DW-SSFP. (d) Example DW-SSFP images acquired with multiple nominal flip-angles at 7T reveal how a change in flip-angle yields changes in SNR, with the impact of the nominal flip angle and B_1_ clearly visible between and within DW-SSFP datasets, consistent with the signal simulation in (b). As the nominal flip angle increases, a bright concentric ring in the DW-SSFP data moves radially towards the edge of the brain. (b) and (d) depict the DW-SSFP signal change (CNR).

When considering B_1_ inhomogeneity, the sensitivity of DW-SSFP to flip angle first leads to spatially varying SNR across the brain (Fig. 1d). Second, a different flip angle also translates into a different “effective b-value” (Tendler et al., 2019) in DW-SSFP. When performing DW-SSFP experiments at 7T, even when incorporating the DW-SSFP signal model (Buxton, 1993), non-Gaussian diffusion (due to restrictions in tissue) can lead to B_1_-dependent diffusivity estimates across the brain. This is analogous to acquiring a dataset with different b-values across the brain with a standard DW-SE experiment.

Given a B_1_ field map, we propose an approach to account for these issues by acquiring a pair of DW-SSFP datasets at two different flip angles. This dual-flip angle approach has two advantages: Firstly, our flip angles can be chosen so different regions of tissue have high SNR in the individual datasets (Foxley et al., 2014b). We can subsequently combine the datasets in a manner to yield high SNR diffusivity estimates over the entire brain. Secondly, we can modify the DW-SSFP signal equation to account for how the measured apparent diffusion coefficient (ADC) varies with flip angle under a simple model of non-Gaussian diffusion. From this, we can explicitly model the relationship between the effective b-value and flip angle (Tendler et al., 2019). Here we describe a method to subsequently derive diffusivity estimates over the entire brain sample interpolated to a single SNR-optimal effective b-value, removing the influence of B_1_.

## Theory

### Dual-flip angle acquisition to optimise diffusion contrast

It is possible to apply a desired flip angle anywhere in the brain given accurate knowledge of the B_1_ distribution and appropriate choice of nominal flip angle. This effect is demonstrated in Fig. 1d, which displays a single slice through a DW-SSFP dataset where the nominal flip angle is changed by 10° increments from 5° to 115°. By changing the nominal flip angle, a bright concentric ring is seen to move radially from the centre of the brain towards the edge.

An arbitrarily optimized flip angle for the DW-SSFP signal can therefore be predictably positioned with knowledge of B_1_. We propose that the signal dependency on B_1_ can be mitigated by acquiring data with an optimized pair of flip angles. Figure 2 outlines our proposed optimization procedure, which aims to produce high contrast across the entire brain. The goal is to identify an optimal pair of nominal flip angles based on the predicted diffusion contrast (here defined as the difference between the non-diffusion and diffusion weighted signals). An ideal flip angle pair would achieve both high and homogeneous contrast over a large range of fractional B_1_ (Fig. 2). To achieve this, DW-SSFP contrast curves were generated for every pair of flip angles (Fig. 2a) and their mean (μ) and standard deviation (σ) over a range of B_1_ values were determined. To identify a flip angle pair that represented a balance of high contrast and homogeneity across a range of B_1_, we calculated the variance-normalized mean (μ/σ) of all flip angle pairs (Fig. 2b), and chose the peak value as our optimal pair of flip angles (Fig. 2c). We considered a range of 30-100% of the maximum B_1_ (Fig. 2a) to ensure that the optimization is not dominated by a minority of voxels with very low B_1_.

**Figure 2:**
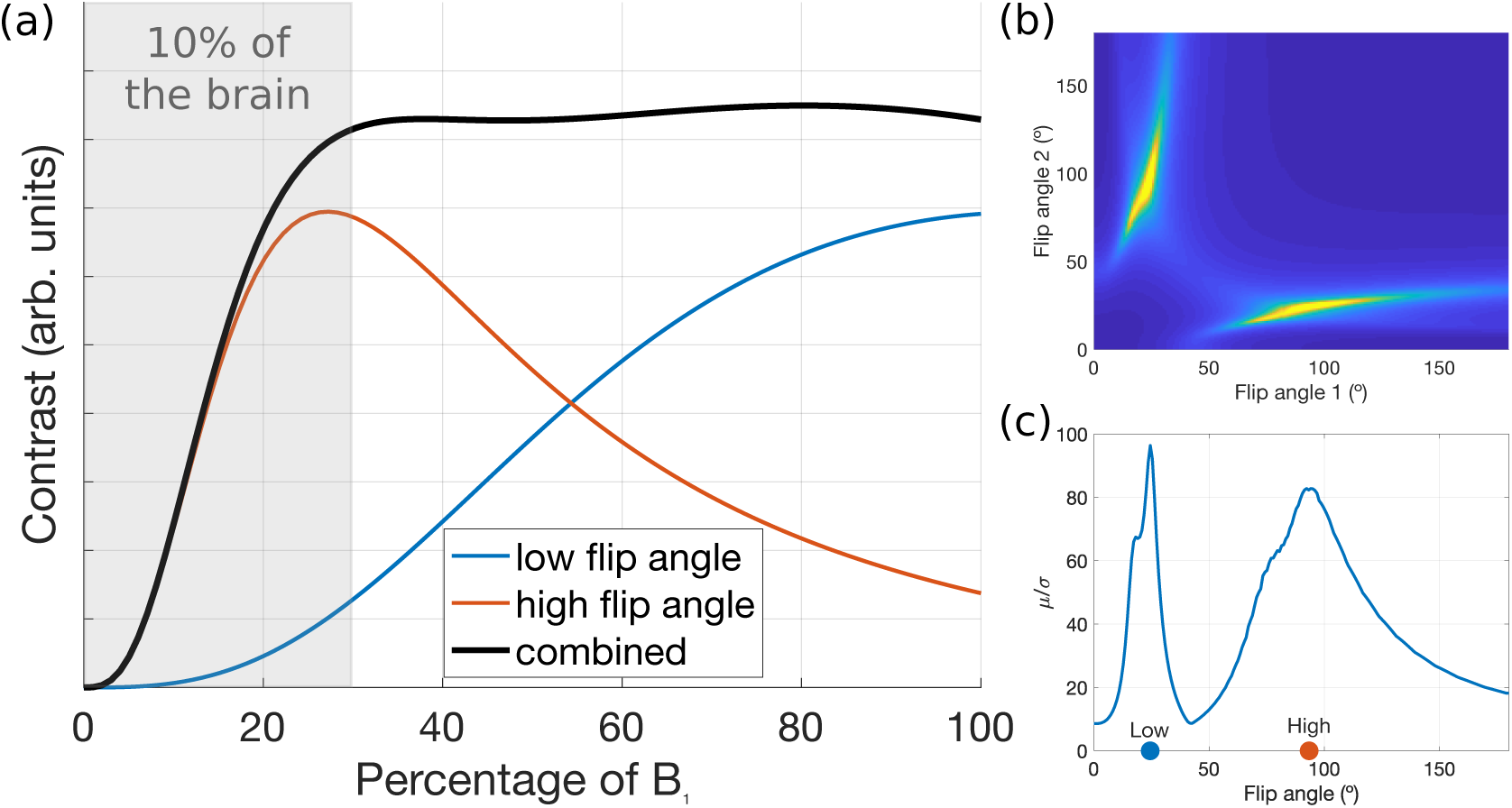
Optimization used for the two-flip angle DW-SSFP acquisition. The red and blue curves in (a) show the variation in diffusion contrast (difference of diffusion weighted and non-diffusion-weighted signal) across the brain as a function of B_1_. We combined the contrast curves across pairs of flip angles to maximize the quantity *μ*/*σ* (b), where *μ* is the mean contrast across the B_1_ range and *σ* is the standard deviation. This metric aims for maximum contrast with minimum variation across the brain (black curve on the left). Here, we only considered a range of 30%-100% of maximum B_1_ for our calculations of *μ* and *σ* which corresponds to ∼90% of the brain, so as not to have the optimization dominated by a minority of brain voxels where contrast changes rapidly with flip angle. Our simulations estimated a CNR-optimal flip angle pair of 24° and 94° (c). Simulation parameters were approximately matched to our protocol at 7T: *T*_1_ = 500 ms, *T*_2_ = 30 ms, ADC = 1 · 10^−4^ mm^2^/s, TR = 30 ms, diffusion gradient amplitude= 52 mT/m, diffusion gradient duration= 14 ms.

### A DW-SSFP effective b-value

In the presence of non-Gaussian diffusion, ADC estimates are highly susceptible to flip-angle variations in DW-SSFP (Tendler et al., 2019). This is a direct result of the signal representing a linear mixture of coherence pathways with different b-values, where the relative weight of pathways is determined by the flip angle. Figures 3a and b show simulations comparing multi-shell DW-SE and multi-flip angle DW-SSFP signal attenuation for systems defined by a single diffusion coefficient (Gaussian) vs a gamma distribution of diffusivities (non-Gaussian). Under Gaussian diffusion, the diffusion attenuation curves (blue lines) do not overlap for different diffusion coefficients. However, for non-Gaussian diffusion (orange lines), the diffusion attenuation curves cross through the blue lines, indicating a change in the measured ADC. The dependence of ADC on b-value in DW-SE is well established. For DW-SSFP a similar effect exists, where the effective b-value is indirectly modified via the flip angle.

**Figure 3:**
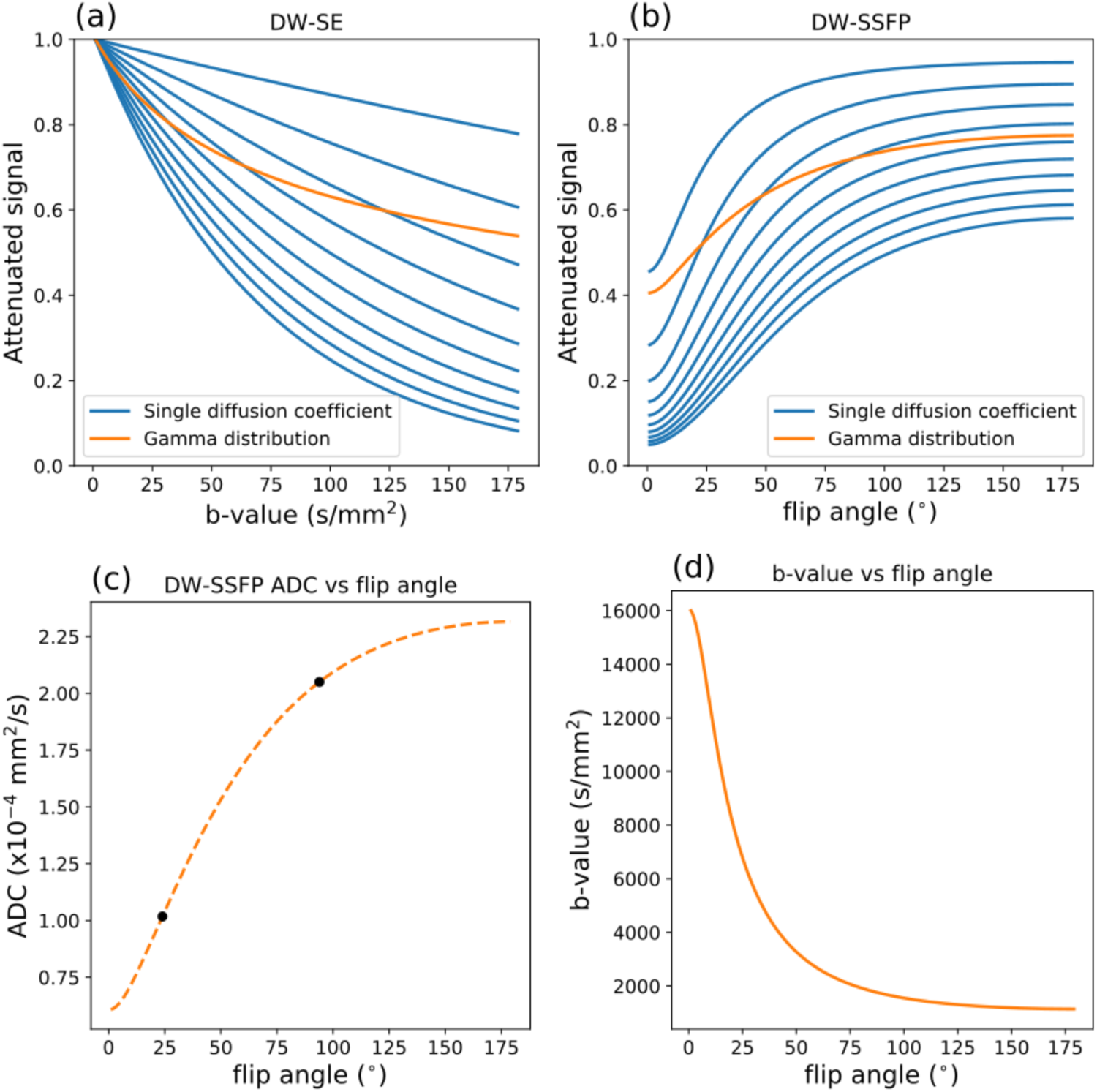
Simulating non-Gaussian diffusion effects on ADC. (a/b) Comparison of the diffusion attenuation of a multi-shell DW-SE (a) and multi-flip angle DW-SSFP (b) experiment in a system defined by a single diffusion coefficient (blue lines) or a gamma distribution of diffusivities (orange line). Non-Gaussian diffusion (gamma distribution of diffusivities) leads to variable measurements of ADC with b-value/flip angle. Given ADC estimates from the DW-SSFP signal at two-flip angles (c – black dots), we can fit a gamma distribution of diffusivities (c – orange dashed line), which predicts the ADC at any given flip angle. (d) As described in (Tendler et al., 2019), this can further be used to describe the system at a well-defined b-value (d).

In DW-SSFP, this flip angle dependence on ADC prevents a simple (weighted) averaging of ADC estimates across acquisitions with multiple flip angles (to produce a composite ADC map with homogeneous SNR). This additionally leads to spatially-dependent ADC estimates within a single DW-SSFP dataset, arising due to B_1_ inhomogeneity combined with non-Gaussian diffusion.

A solution to this problem may be as follows: From DW-SSFP data in a voxel acquired at two nominal flip angles, we can use the standard Buxton signal model (Buxton, 1993) to calculate an ADC estimate for each flip angle (Fig. 3c – black dots). We can then fit these ADC estimates with a modified Buxton model that incorporates non-Gaussian diffusion (Fig. 3c – orange line). Based on this characterisation of the non-Gaussianity in our voxel, we can calculate the ADC estimate at any given flip angle by interpolating (or extrapolating) along the DW-SSFP signal model curve. Thus, we can eliminate the influence of B_1_ on our ADC estimates by calculating the ADC at the same flip angle over the entire brain.

In DW-SSFP, parameters in addition to the flip angle can lead to changes in the estimated ADC, including the relaxation times T_1_ and T_2_. Motivated by this, recent work (Tendler et al., 2019) redefined the DW-SSFP signal in terms of an ‘effective’ b-value, b_eff_, the equivalent b-value with the DW-SE sequence that leads to the same estimate of ADC (Fig. 3d) under an identical model of non-Gaussian diffusion. This definition accounts for the influence of these different parameters on the measured ADC, in contrast to previous work (Miller et al., 2012), which defined b_eff_ in terms of the DW-SSFP diffusion attenuation. The new definition of b_eff_ allows for direct comparisons of results to equivalent DW-SE data assuming the same model of non-Gaussianity. The effective b-value can be additionally chosen to account for the variable SNR of DW-SSFP data at different nominal flip angles/B_1_, to optimize the SNR of the ADC estimates over the entire brain (described in Supplementary Material).

In this work, we utilise a Gamma distribution of diffusivities to capture non-Gaussian diffusion (Jbabdi et al., 2012; Tendler et al., 2019). We chose a Gamma distribution in part because it only adds a single extra free parameter in comparison to a Gaussian diffusion model. A more common model of non-Gaussianity is bi-exponential diffusion, but this model is both relatively crude and requires the addition of two free parameters. Furthermore, the gamma distribution is only defined for positive diffusion coefficients and can be parameterised in terms of a mean, *D*_*m*_, and standard deviation, *D*_*s*_. This allows for the incorporation and correction for the non-Gaussian diffusion properties of DW-SSFP data acquired at only two-flip angles. Further details of this framework can be found in (Tendler et al., 2019).

## Methods

### Sample preparation

Data were acquired in post-mortem human brains (n=5), comprised of two control brains and three brains from patients diagnosed with amyotrophic lateral sclerosis (ALS). Brains were extracted from the skull within 72 hours after death. All brains were fixed for at least 45 days prior to scanning, with four brains fixed in 10% PBS buffered formalin and one brain fixed in 10% formalin. Prior to scanning, brains were removed from formalin and submerged in a perfluorocarbon liquid (Fluorinert FC-3283, 3M). The study was conducted under the Oxford Brain Bank’s generic Research Ethics Committee approval (15/SC/0639).

### MRI Data acquisition protocol

Data were obtained over the entire brain of each post-mortem sample on a human 7T Siemens whole body scanner (32ch-receive/1ch-transmit head coil). For each brain, DW-SSFP datasets were acquired at two-flip angles (24° and 94°), chosen based on the optimization described above. At each flip angle, 120 diffusion directions (q = 300cm^-1^) and six non-diffusion weighted datasets were acquired (resolution = 0.85·0.85·0.85 mm^3^), with the same set of directions for both flip angles. To prevent banding artefacts in the non-diffusion weighted datasets, a slight diffusion gradient was applied along (x,y,z) = (0.557,0.577,0.577) to serve as a spoiler (q = 20cm^−1^) (Zur et al., 1988).

To aid in DW-SSFP quantification, we also acquired: B_1_ maps with an actual flip angle (AFI) acquisition (Yarnykh, 2007); T_1_ maps from a turbo inversion-recovery (TIR) sequence; and T_2_ maps from a turbo spin-echo (TSE) sequence. Full details of the acquisition protocol are provided in Table 1.

**Table 1:** MRI imaging parameters. The imaging parameters of the DW-SSFP dependency acquisitions (AFI, TIR and TSE) are representative of the parameters used, small modifications were made to these acquisitions as protocols evolved.

| <u>DW-SSFP</u> |  | <u>Turbo inversion-recovery (TIR)</u> |  |
| --- | --- | --- | --- |
| q-value ($\text{cm}^{-1}$) | 300 | Resolution ($\text{mm}^3$) | 0.9·0.9·0.9 |
| Diffusion Gradient Duration (ms) | 13.56 | Number of inversions | 6 |
| Diffusion Gradient Strength ($\text{mTm}^{-1}$) | 52 | TE (ms) | 14 |
| Flip angles ($^\circ$) | 24 and 94 | TR (ms) | 1000 |
| No. directions (per flip angle) | 120 | TIs (ms) | 30, 60, 120, 240, 480 & 935 |
| No. non-DW (per flip angle) | 6 ($q=20\text{ cm}^{-1}$) | Flip angle ($^\circ$) | 180 |
| Resolution ($\text{mm}^3$) | 0.85·0.85·0.85 | GRAPPA acc. factor | 3 |
| TE (ms) | 21 | Bandwidth (Hz per pixel) | 130 |
| TR (ms) | 28 | Acquisition time (per TI) | 40:49 |
| EPI factor | 3 | Number of averages | 1 |
| Bandwidth (Hz per pixel) | 393 |  |  |
| Acquisition time (per direction/non-DW) | 5:47 | <u>Turbo spin-echo (TSE) – $T_2$</u> |  |
| Acquisition time (per flip angle) | 12:08:42 | Resolution ($\text{mm}^3$) | 0.9·0.9·0.9 |
| No. of averages | 1 | Number of echoes | 6 |
|  |  | TEs (ms) | 13, 25, 38, 50, 63 & 76 |
| <u>Actual flip-angle imaging (AFI) – $B_1$</u> |  | TR (ms) | 1000 |
| Resolution ($\text{mm}^3$) | 3·3·3 | Flip angle ($^\circ$) | 180 |
| TE (ms) | 1.5 | GRAPPA acc. factor | 2 |
| TR<sub>1</sub>/TR<sub>2</sub> (ms) | 4.4/11 | Bandwidth (Hz per pixel) | 166 |
| Flip angle ($^\circ$) | 60 | Acquisition time (per TE) | 36:01 |
| Bandwidth (Hz per pixel) | 630 | Number of averages | 1 |
| Acquisition time | 0:41 |  |  |
| Number of averages | 1 |  |  |

### Data Processing

All coregistrations between and within imaging modalities were performed with a 6 degrees-of-freedom (translations and rotations) co-registration via FLIRT (Jenkinson et al., 2002; Jenkinson and Smith, 2001). A Gibbs ringing correction was performed on the DW-SSFP, TIR and TSE datasets (Kellner et al., 2016). T_1_ and T_2_ maps were generated from the TIR and TSE data via a voxelwise fit assuming mono-exponential signal evolution. B_1_ maps were generated from the AFI datasets via the processing outlined in the original publication (Yarnykh, 2007) All data were processed and analyzed using the FMRIB software library (FSL) (Jenkinson et al., 2012) and Python (Millman and Aivazis, 2011). A diffusion tensor model that incorporates the full DW-SSFP Buxton signal model, including T_1_, T_2_ and B_1_ (Buxton, 1993) was fitted to the DW-SSFP data using cuDIMOT (Hernandez-Fernandez et al., 2019).

This work incorporates two versions of the diffusion tensor model, one which fits DW-SSFP data acquired at one-flip angle and one that fits a tensor to data at two-flip angles simultaneously. The latter analysis outputs a unique set of eigenvalues, *L*_1,2,3_, for the DW-SSFP data acquired at each flip angle, but is constrained to a shared set of eigenvectors, 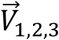. All comparative analyses were done solely over white matter, with white matter masks generated using FAST (Zhang et al., 2001), followed by manual removal of any remaining grey matter regions.

### Comparison of PDD estimates acquired with one- and two-flip angle acquisitions

To compare the resulting diffusion eigenvectors between the one- and two-flip angle acquisitions, a time-matched comparison was performed. A subset of the data (60 directions at each flip angle) were selected and fitted with the two-flip angle DW-SSFP tensor model described above. These model fits were compared to the results obtained from fitting to all 120 directions of DW-SSFP data acquired at one-flip angle only. The subset of directions was chosen for maximally even coverage in the angular domain, ensuring a fair comparison of an equal number of directions and similar angular resolution between the one- and two-flip angle analyses.

The one-vs two-flip angle PDD estimates were compared using a measure of angular uncertainty from the orientations of samples from the posterior distribution (Jbabdi et al., 2012). The resulting estimate (defined as a scalar between 0 and 1, where a larger number corresponds to a higher uncertainty) reflects the extent to which tractography can be successfully performed within the brain.

### Combination of eigenvalue estimates at two-flip angles to a single effective b-value

In order to estimate voxelwise ADC maps for a single b_eff_ across the brain, we first need to fit the parameters of our model of non-Gaussian diffusion. As in our previous work (Tendler et al., 2019), we use a gamma distribution of diffusivities with mean, *D*_*m*_ and standard deviation, *D*_*s*_ in each voxel. Below we describe a robust procedure to achieve this.

For each voxel, we first obtain ADC estimates obtained separately at a low and high flip angle using the full Buxton model (Buxton, 1993). These ADC estimates are fit with simulated ADCs for a given set of *D*_*m*_ and *D*_*s*_ (with a measured T_1_ and T_2_) as follows:

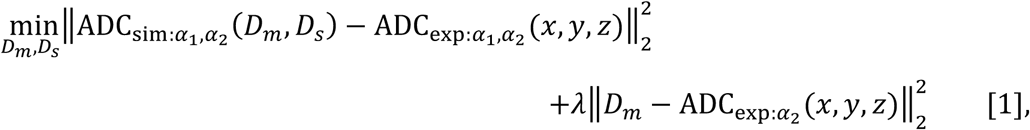

where *α*_1_ and *α*_2_ are the voxelwise DW-SSFP flip angles (defined as the nominal flip angles scaled by the B_1_ gain factor), 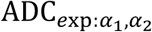 are the voxelwise experimental ADC estimates at each flip angle and 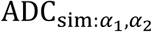 are the simulated ADC estimates for a given *D*_*m*_ and *D*_*s*_ at each flip angle. To prevent overfitting to the experimental data, a regularisation term can be additionally introduced to ensure the estimate of *D*_*m*_ remains on the order of the ADCs estimated separately from the one-flip angle analyses (*λ* is the regularisation parameter). Details of the incorporation of a Gamma distribution of diffusivities into the Buxton model, which form the definition of ADC_sim_(*D*_*m*_,*D*_*s*_) can be found in (Tendler et al., 2019).

Here, tensor estimates were first obtained using the full set of 120 DW-SSFP directions obtained at each flip angle. Fitting a tensor model to the experimental data, a shared set of 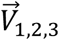 and unique *L*_1,2,3_ at each flip angle were estimated separately. In the second stage, the eigenvectors were then fixed and the posterior distribution of the *L*_1,2,3_ estimates were fit using Eq. [1] to determine voxelwise estimates of *D*_*m*_ and *D*_*s*_ as described (Tendler et al., 2019). Fitting was performed separately for each eigenvalue to determine a unique *D*_*m*_ and *D*_*s*_ for *L*_1_, *L*_2_ and *L*_3_. Fitting was performed in Python using SciPy *curve_fit*, implemented with the Levenberg-Marquardt algorithm (Levenberg, 1944) and accelerated using the Numba compiler (Lam et al., 2015).

*L*_1,2,3_ estimates were subsequently derived over the entire brain in terms of a single b_eff_ (Fig. 3d) using the framework in (Tendler et al., 2019) (described in Supplementary Material Fig. S1). The b_eff_ was chosen to account for the variable SNR of the *L*_1,2,3_ estimates over the entire brain to produce SNR-optimal results, as described in the Supplementary Material.

## Results

### Comparison of PDD estimates acquired with one- and two-flip angle acquisitions

The benefit of the time-matched two-flip angle approach for overcoming B_1_ dependent CNR in PDD estimates is illustrated in Fig. 4. PDD estimates derived from data acquired with a 24° nominal flip angle (120 directions) display greater coherence between voxels near the centre of the brain (Fig. 4 orange box – 24°). As the scanner sets the nominal flip angle of 24° to be matched to this region, we expect the CNR to be maximized (as predicted in Fig. 1b). Within this region, clear delineation of the striations within the internal capsule are visible. In this same region, the PDD estimates with a 94° nominal flip angle (120 directions) are less coherent (Fig. 4 orange box – 94°). At the brain boundary where the actual flip angle is far below the nominal flip angle, the opposite is true. The PDD estimates at 94° reveal clear depiction of cortical folding patterns (Fig. 4 red box - 94°), which are corrupted by noise at 24° (Fig. 4 red box - 24°). In comparison, PDD estimates of the two-flip angle data (120 directions, 60 directions at 24° and 60 directions at 94°) (Fig. 4 24°+94°) demonstrate that regionally dependent benefits associated with each single-flip analysis are captured by the two-flip angle approach. In this combined scan time-matched dataset, it is possible to visualize cortical folding, whilst maintaining the striations within the internal capsule.

**Figure 4:**
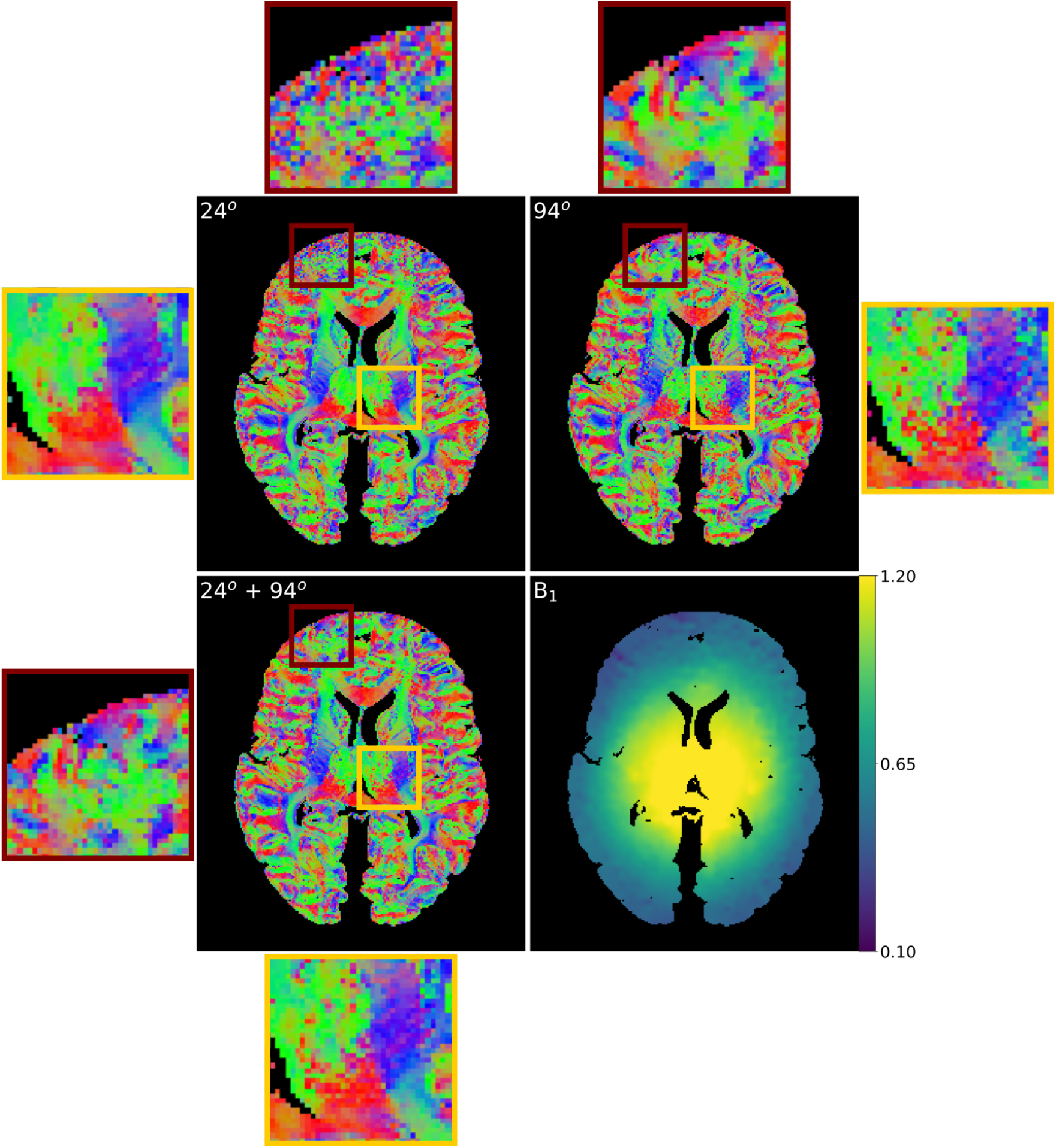
Visual comparison of the PDD estimates. For the 24° dataset, B_1_ inhomogeneity leads to incoherent PDD estimates near the brain boundary (red box), with coherent PDD estimates near the centre of the brain (orange box). For the 94° dataset, the converse is true. By fitting with two-flip angles (24° + 94°), we obtain a good compromise between the low and high flip angle datasets, yielding coherent PDD estimates over the entire brain.

Figure 5 shows how the angular uncertainty varies as a function of B_1_, where low uncertainty indicates high CNR. In all five datasets, the low B_1_ near the periphery of the brain leads to a higher angular uncertainly in the 24° datasets when compared to those acquired at 94o. In areas of high B_1_ the opposite is true, in agreement with Fig. 4. The dual-flip approach (24°+94°) is able to generate PDD estimates with angular uncertainty closer to the best performance obtained for the one-flip angle datasets at the extremes of high or low B_1_, and in many cases outperforms either single-flip dataset between these values (i.e. where the curves cross in Fig. 5). A histogram (Fig. 5, bottom right) shows the broad range of B_1_ values sampled in our post-mortem brains.

**Figure 5:**
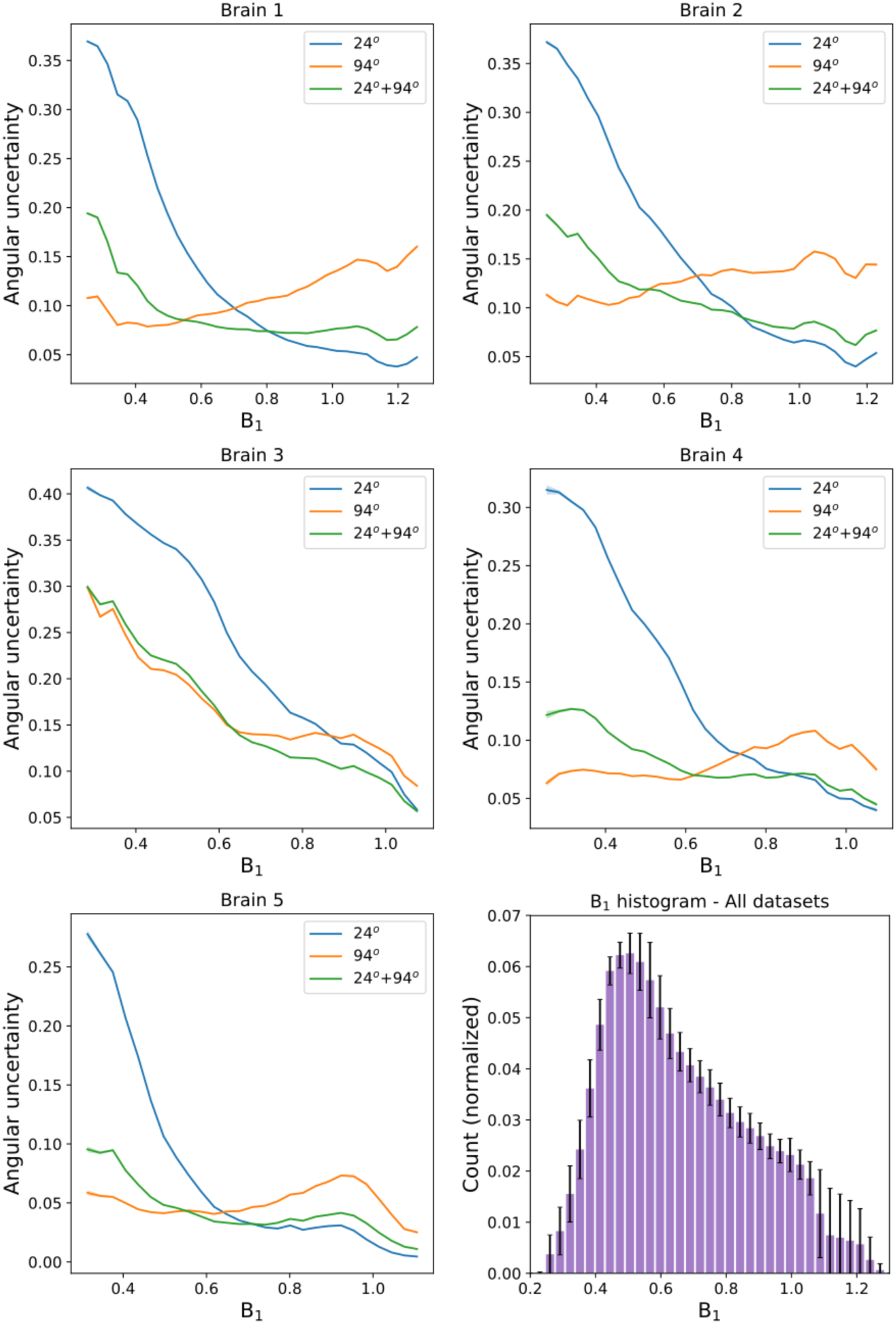
PDD angular uncertainty as a function of B_1_. In all 5 brains, it can be seen that PDD angular uncertainty estimates are reduced in areas of low/high B_1_ for the 94°/24° datasets respectively. After the proposed combination of two-flip angle data (24° + 94°), the PDD uncertainty estimates are closer to those of the single-flip angle within their respective regions of high CNR. Plots generated in white matter only from the PDD uncertainty and B_1_ maps for each of the five datasets. The standard error of PDD dispersion values are plotted for each brain, but due to the large number of points per bin these error bars are too small to be visualized. The B_1_ histogram (bottom right) reveals that the B_1_ values sampled within these datasets spans a wide range of B_1_, with error bars denoting the standard deviation over the five datasets.

Figure 6 shows a map of the difference in uncertainty between the one- and two-flip angle results. While there are parts of the brain where acquisition at a single, optimal flip angle provides slightly lower uncertainty compared to the two-flip angle approach (light red), over the entire dataset the dual-flip approach provides a net gain (dark blue). By creating a histogram of the difference in PDD angular uncertainty between the one- and two-flip angle analyses (Fig. 7), we can see an increased fraction of voxels with the two-flip angle approach that have a large reduction in uncertainty in comparison to 24° all brains) and 94° (4/5 brains) (blue curves above red). The opposite is true for small differences in angular uncertainty (red curves above blue). The overall improvements in angular uncertainty for the two-flip angle approach vs 94° are reduced in comparison to 24°, reflecting the large number of voxels at 24° which have high angular uncertainty (Fig. 5).

**Figure 6:**
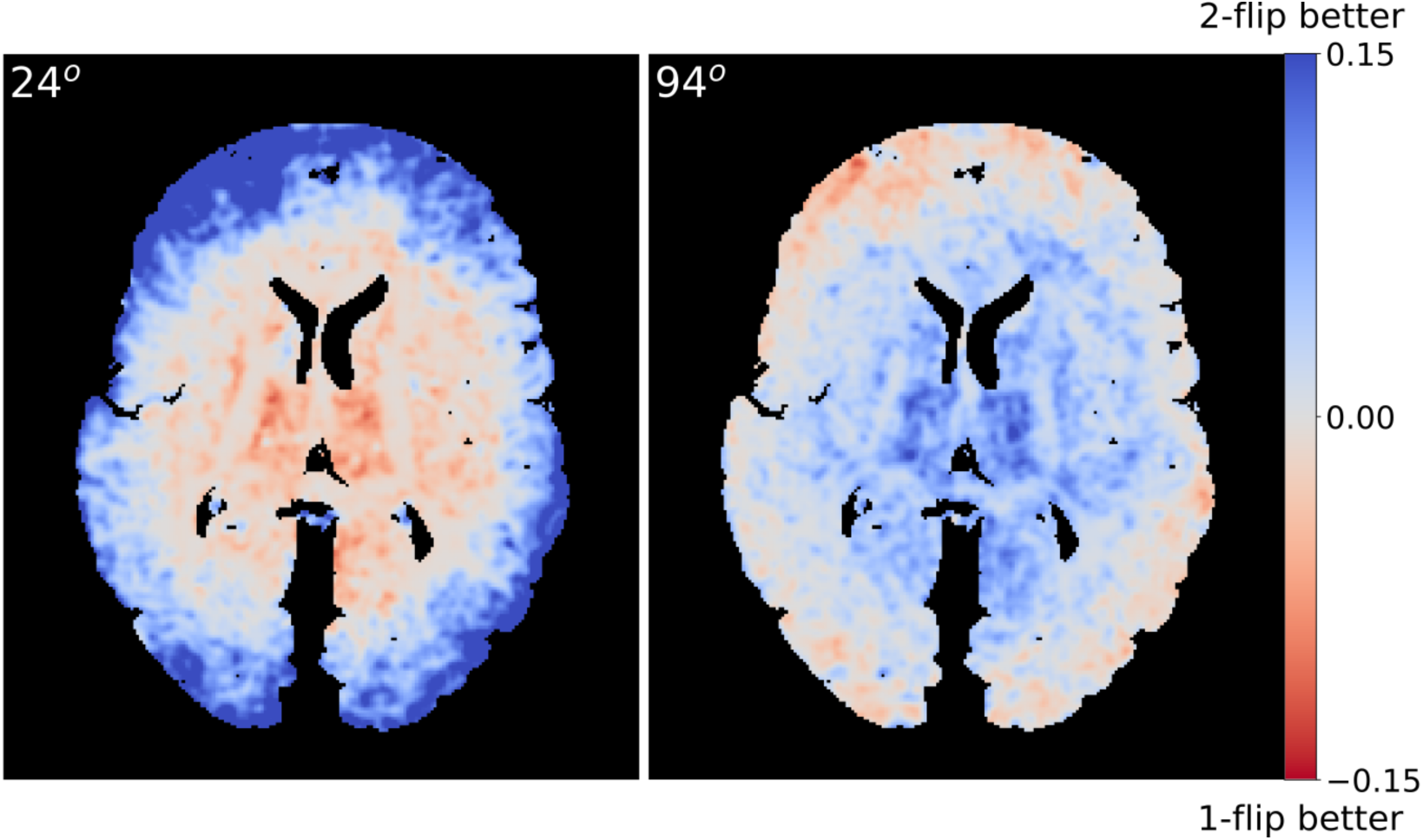
Visual comparison of the differences in PDD angular uncertainty. Positive values (blue) display regions where the two-flip angle approach outperforms the single-flip angle, whereas negative values (red) display the opposite. Areas of higher/lower uncertainty are in good visual agreement with the coherence of the PDD estimates in Fig. 4. To aid visualization, the uncertainty differences were smoothed with a Gaussian filter (standard deviation = 1 mm).

**Figure 7:**
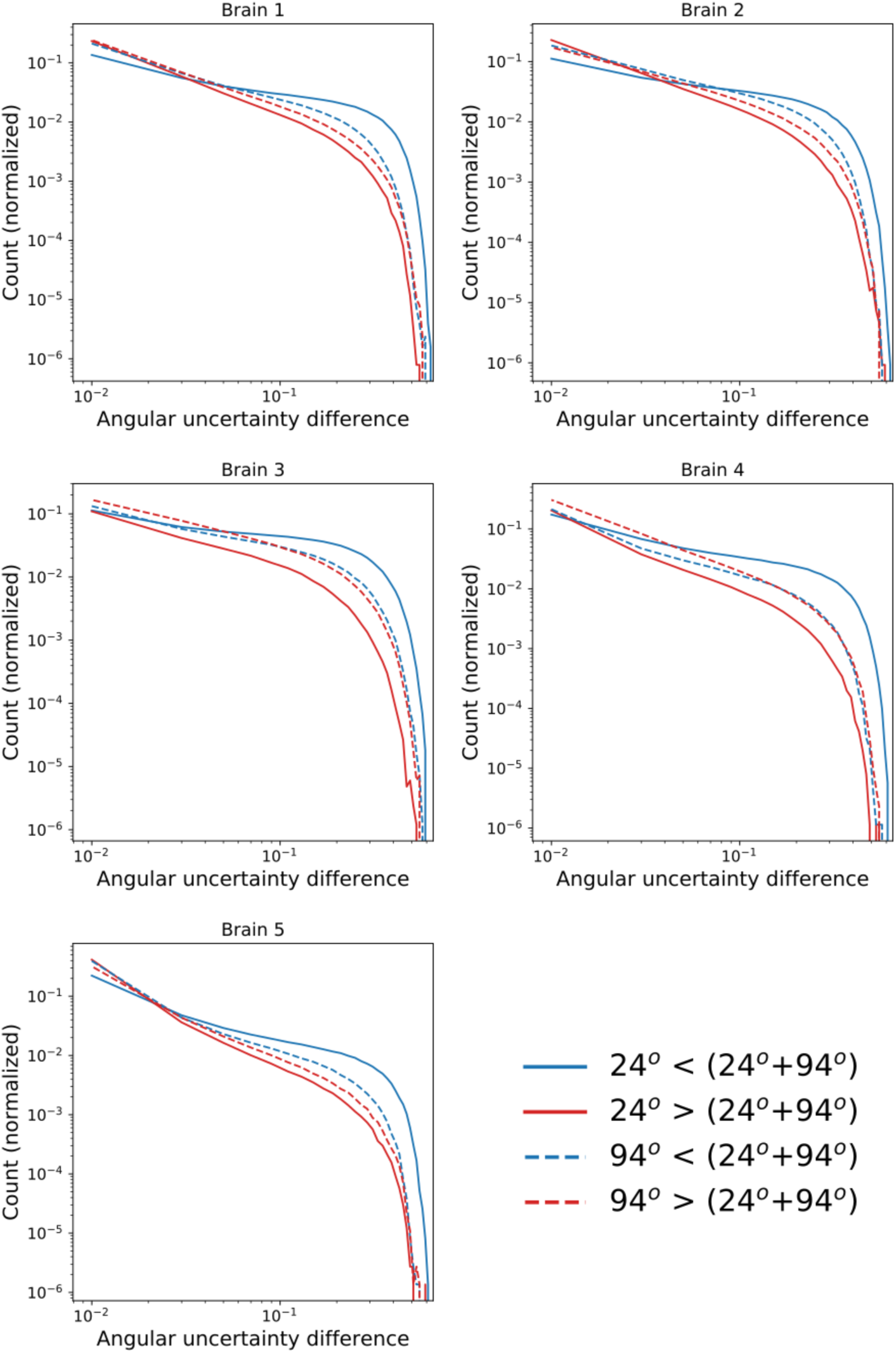
Quantitative comparison of the differences in PDD angular uncertainty. These PDD uncertainty difference histograms represent the number of voxels where the one-/two-flip angle PDD estimates outperforms the other. Here, solid/dashed lines refer to the difference between the 24°/94° and the two-flip angle approach respectively. Blue lines indicate the number of voxels that the two-flip angle approach outperforms the single-flip angle, whereas the red lines display the opposite. A log scale is used on both the x- and y-axes.

### Combination of eigenvalue estimates at two-flip angles to a single b_eff_

*L*_1,2,3_ estimates calculated from DW-SSFP data at 24° and 94° (Fig. 8) display observable differences in the derived diffusivity values, overall showing an increased diffusivity estimate at 94° (confirmed in Fig. 9). Previous work (Tendler et al., 2019) makes clear that effective b-values are overall higher with lower flip angles, which would be consistent with these variations in diffusivity being driven by restriction in tissue. Furthermore, this indicates that we cannot simply average the eigenvalue estimates acquired at different DW-SSFP flip angles, as it would combine maps with distinct ADC estimates at each flip angle. *L*_1,2,3_ maps at the SNR-optimal b_eff_ (determined as b_eff_ = 7600 s/mm^2^ – details of derivation in Supplementary Material) show a reduced inhomogeneity in comparison to the 24° dataset, and improved SNR when compared to 94°.

**Figure 8:**
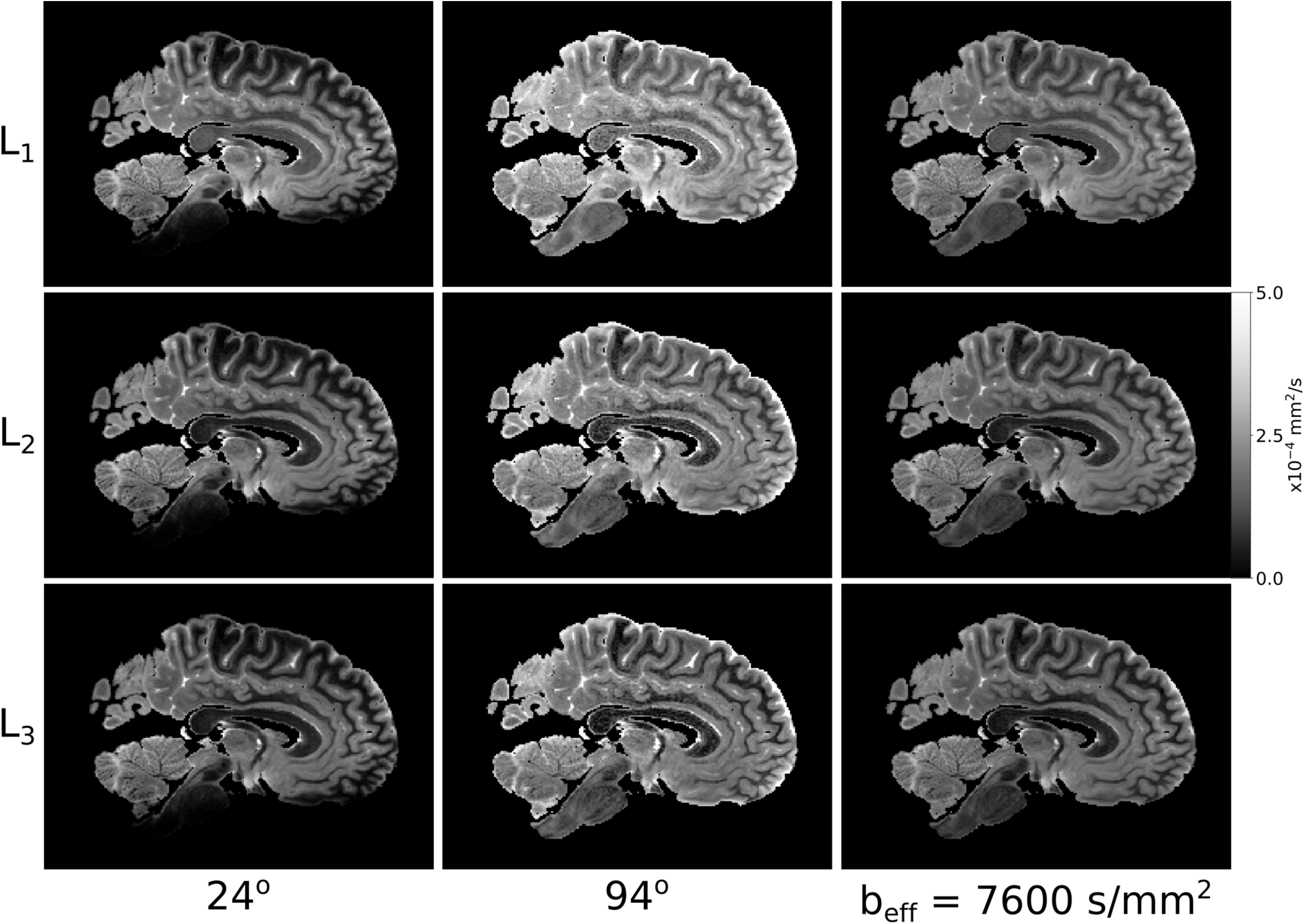
Visual comparison of the *L*_1,2,3_ estimates. Differences in the *L*_1,2,3_ maps at 24° and 94° agrees with the expectation that within a non-Gaussian regime, increased flip angle in DW-SSFP yields higher diffusivity estimates (Fig. 3c). The *L*_1,2,3_ maps at b_eff_ = 7600 s/mm^2^ reveal improved SNR vs 94° and more homogenous diffusivity estimates over tissue vs 24°.

**Figure 9:**
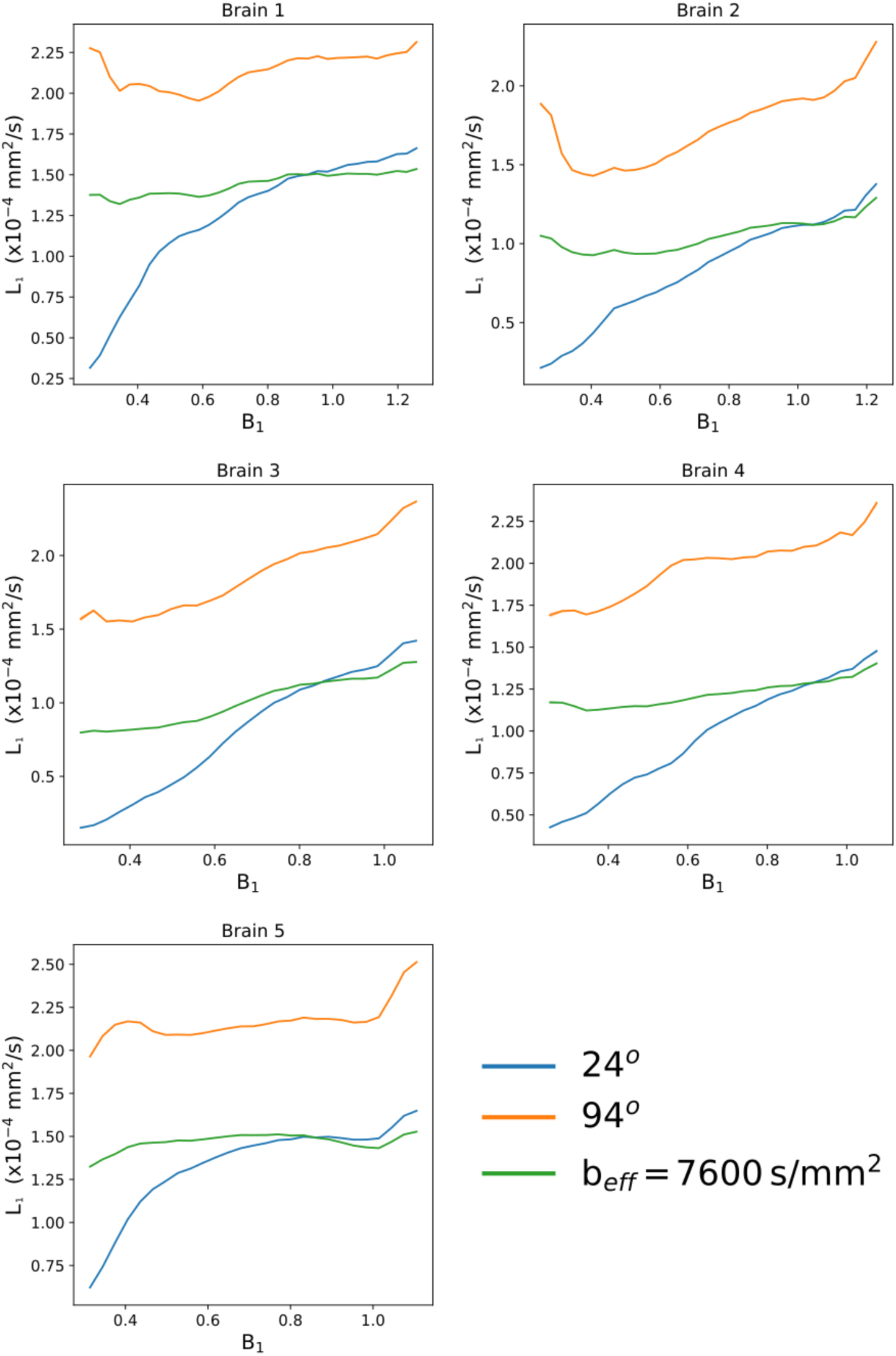
Quantitative comparison of *L*_1_ estimates vs B_1_. Here we observe an increased *L*_1_ estimate in DW-SSFP data acquired at 94°, in agreement with (Tendler et al., 2019) and Fig. 3c. The *L*_1_ estimates at b_eff_ = 7600 s/mm^2^ display a flatter distribution, consistent with removal of the influence of B_1_. Plots generated in white matter only from the *L*_1_ and B_1_ maps for each of the five datasets. The standard error of *L*_1_ estimates within each bin are plotted for each brain, but due to the large number of points per bin these error bars are too small to be visualized.

As shown in Fig. 9, the reconstructed *L*_1_ estimates at b_eff_ = 7600 s/mm^2^ give good agreement to the 24o results at high B_1_, whilst maintaining a flatter distribution at lower B_1_ within all five brains. The crossing point of the *L*_1_ curves at 24° and b_eff_ = 7600 s/mm^2^ reveals the approximate flip angle along *L*_1_ where b_eff_ = 7600 s/mm^2^.

Fractional anisotropy (FA) maps over all five brains (Fig. 10) additionally display differences in the estimated FA at 24° and 94° (confirmed in Fig. 11), consistent with restriction along *L*_1,2,3_. These FA maps have an increased sensitivity to noise in comparison to the *L*_1,2,3_ estimates and the FA maps derived from DW-SSFP data at 24°/94° have low SNR at the edge/centre of the brain respectively, consistent with the PDD results in Fig. 4. The FA maps generated at b_eff_ = 7600 s/mm^2^ do not reveal the same spatial variation, yielding high SNR across the brain. The impact of B_1_ is displayed in Fig. 11.

**Figure 10:**
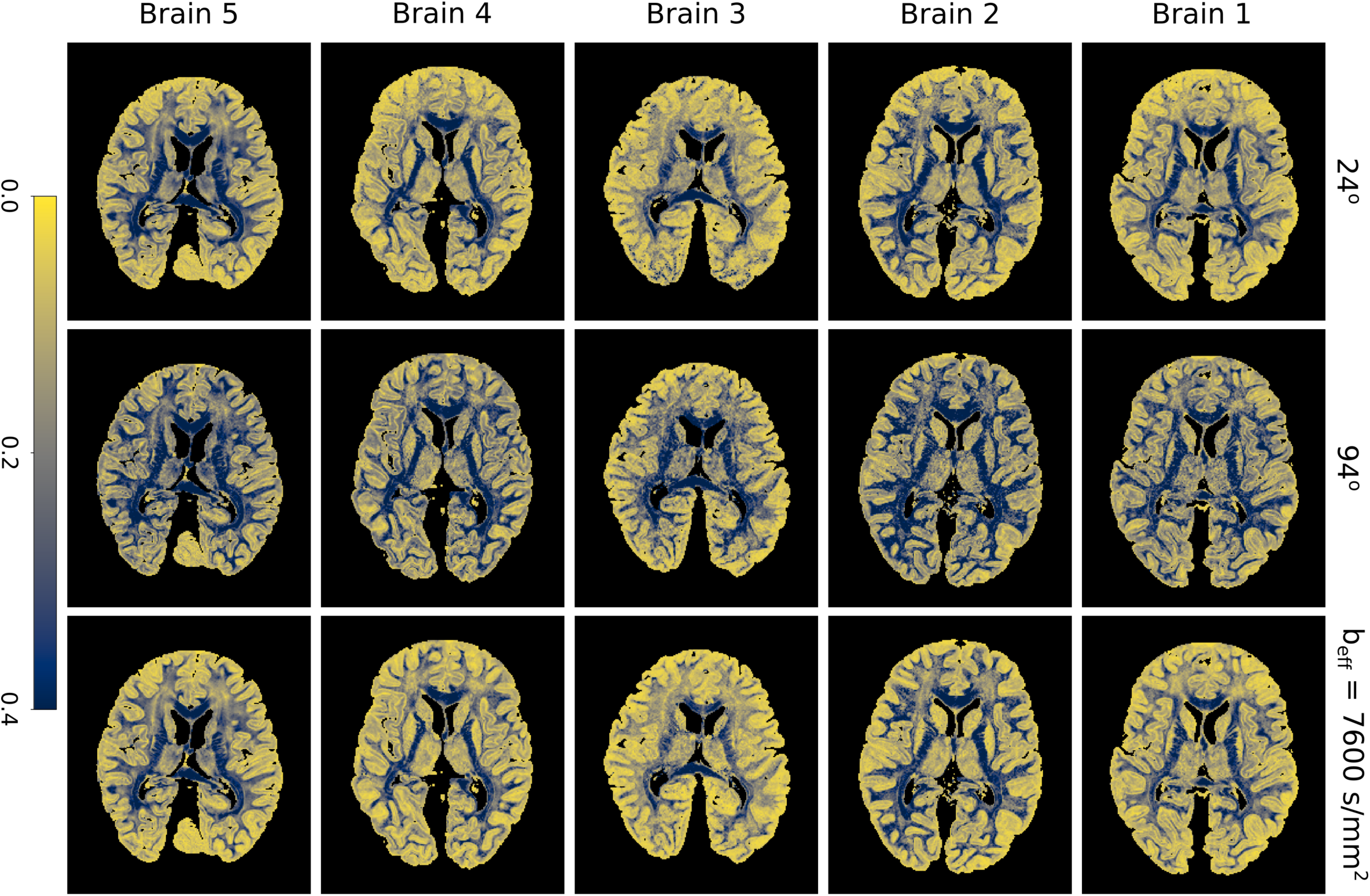
Visual comparison of FA. Differences between the FA maps at 24° and 94° are consistent with differences in non-Gaussianity along the three tensor eigenvectors, additionally revealing a reduced SNR in the derived FA maps near the boundary/centre at 24°/94° respectively. The FA maps at b_eff_ = 7600 s/mm^2^ yield more consistent SNR across the tissue. Colormap chosen to highlight the variable contrast and noise over the brain.

**Figure 11:**
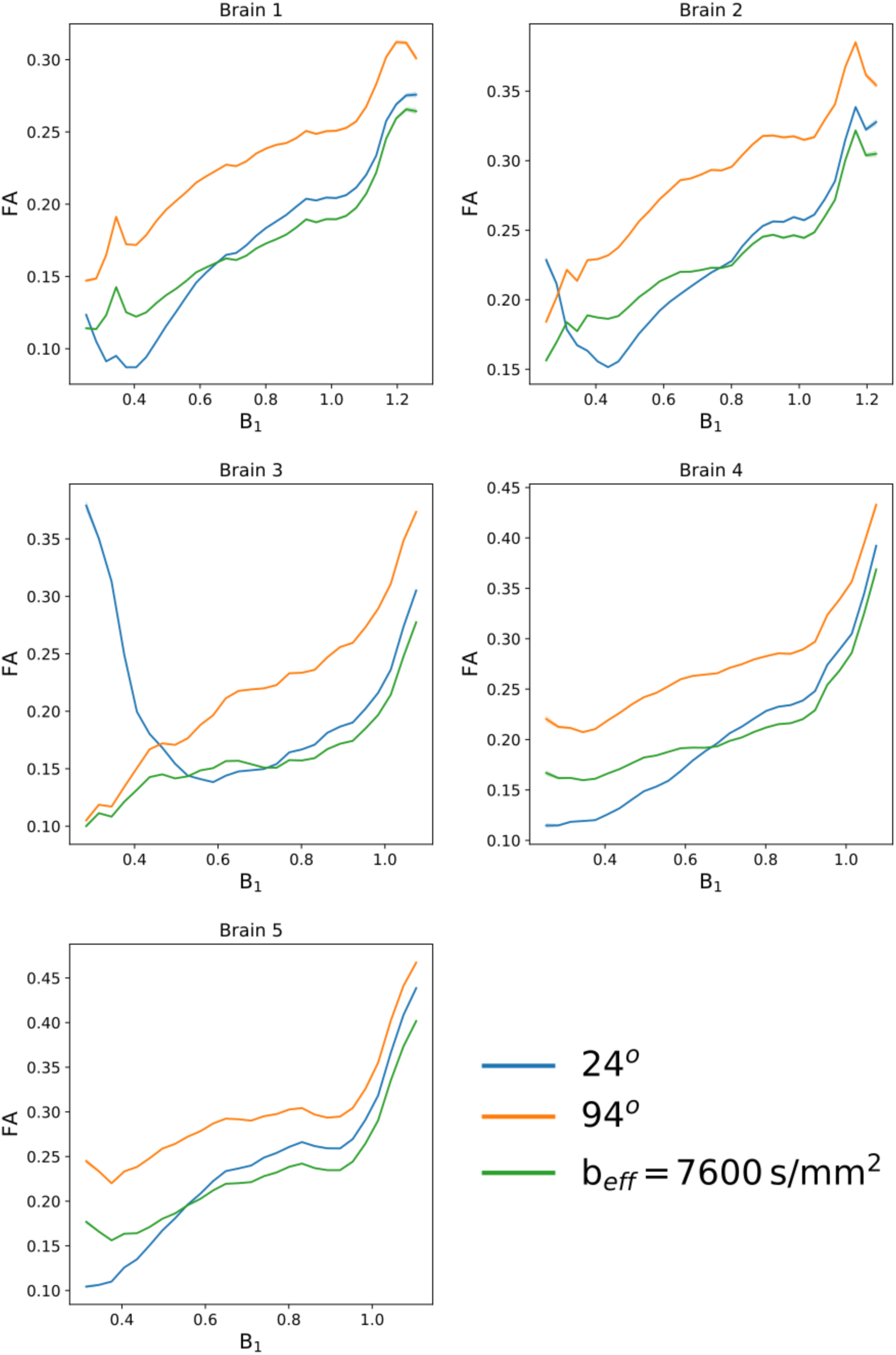
Quantitative comparison of FA estimates vs B_1_. Here we observe an increased FA estimate in DW-SSFP data acquired at 94°, consistent with variations in the non-Gaussian properties of tissue along the estimated eigenvalues. Plots generated in white matter only from the FA and B_1_ maps for each of the five datasets. The standard error of FA estimates within each bin are plotted for each brain, but due to the large number of points per bin these error bars are too small to be visualized.

## Discussion

This work demonstrates how the effects of B_1_ inhomogeneity in DW-SSFP can be accounted for by using data acquired at two-flip angles and an appropriate signal model that captures non-Gaussian diffusion. By utilizing a pair of prescribed flip angles that optimize CNR across a range of B_1_, we provide a means to obtain a homogeneous and interpretable characterization of diffusion across the brain. We demonstrate the potential of this approach by quantifying the spatial profile of angular uncertainty in PDD estimates and diffusivity estimates as a function of B_1_.

Previous work (Foxley et al., 2014a) demonstrated that with a one-flip angle DW-SSFP acquisition, angular uncertainty in PDD estimates was reduced by increasing field strength from 3T to 7T, providing motivation to move to higher field when performing tractography. This reduction in uncertainty would be expected in local regions of tissue due to the higher SNR associated with an increase in field strength, but would be mitigated by the B_1_ effects considered in this work (Fig. 4). Using the two-flip approach described in this paper, PDD estimates at 7T can be obtained over whole post-mortem brain samples (Fig. 4), reducing the number of voxels with high angular uncertainty in tissue regions that experience a sub-optimal flip angle (Fig. 7). Given the pattern of B_1_ and the need for high quality data in central white matter for tractography, there is a particular benefit for tractography into the grey matter. This is a potentially important improvement as such measurements would allow for resolving inter-cortical tracts such as U-fibers as well as more accurately depicting white matter penetration of cortical grey matter away from the gyral crown.

For these post-mortem brain samples, SNR-optimal estimates are predicted to be achieved at a low flip angles. Our SNR-optimal b_eff_ corresponds to an approximate flip angle of 20° − 24° (Supplementary Material Fig. S2d), achieved at B_1_ values of 0.83-1/0.19-0.26 for the 24°/94° datasets. The plots in Fig. 5 show that the two-flip angle approach achieves an angular uncertainty estimate closer to the single flip angle approach in these B_1_ regions and often performs better between these B_1_ values. Further improvement could be achieved by incorporating weighting into the two-flip angle DWSSP tensor model fitting (i.e. weighted least squares), reducing the influence of the DW-SSFP flip angle with high angular uncertainty. This would be particularly noticeable for the 94° case, where at present the high angular uncertainty associated with the 24° datasets near the brain boundary reduces the performance of the two-flip angle method.

An increased estimate of ADC at higher flip-angles (Figs. 8 and 9) demonstrates deviations of the DW-SSFP signal from the Buxton model, consistent with a model of restriction and the results in (Tendler et al., 2019). Our correction reduces the variation of ADC with B_1_ (Fig. 9), in addition to modifying the distribution of derived metrics such as FA (Fig. 11). This allows for more accurate comparisons of diffusivity estimates within different brain regions. Furthermore, as the B_1_ distribution is not reliably calibrated at scan time, our approach allows for comparison of diffusivity estimates between different post-mortem brain samples. The divergence of the 24° and b_eff_ = 7600 s/mm^2^ plots (Figs. 9), emphasizes the influence of B_1_ on measured ADC.

The FA maps in Fig. 10 reveal the trend of reduced SNR at 24° /94° near the centre/edge of the brain, consistent with the PDD (Fig. 4) maps. However, in the eigenvalue estimates at 24°, we observe a sharply decreasing diffusivity estimate in areas associated with very low B_1_ (Figs. 8 and 9), with a distinctive shading near the brain boundary, most notable in the *L*_1_ map. This shading is hypothesised to be additionally driven by the noise floor on our DW-SSFP data, leading to a reduced diffusivity estimate in areas of low signal (Jones and Basser, 2004). Future work will investigate the use of a noise floor correction to account for this bias.

This study was motivated by the interest in understanding whether diffusivity could provide biomarkers that are related to neuropathology in ALS. This necessitates measures of diffusivity in post-mortem tissue that can be compared to histopathological stains. To be meaningful, these diffusivity measures need to be driven primarily by the underlying tissue (as reflected in restrictions that cause non-Gaussian behaviour) rather than confounds like B_1_ inhomogeneity. For example, neurodegenerative diseases such as ALS have been shown to reduce FA *in vivo* (Agosta et al., 2010). A more consistent measurement of FA across white matter, obtained from results at a single b_eff_ (Fig. 10) would allow for more accurate measurements in post-mortem data to corroborate *in vivo* findings. Future work that directly compares diffusivity to histology will consider whether there is evidence for a neuropathological signature in diffusion MRI.

## Conclusion

DW-SSFP at 7T has the potential to provide high signal and contrast diffusion weighted imaging in post-mortem tissue. However, B_1_ inhomogeneity coupled with the dependence of diffusion contrast on flip angle means that the resulting signal is not straightforward to interpret. We proposed to use a multi-flip angle DW-SSFP acquisition alongside a non-Gaussian signal model to account for B_1_ inhomogeneity at 7T. With this method, we can obtain improved estimates of diffusion properties within tissue, including both quantitative diffusivities and fibre orientations.

## Acknowledgements

This study was funded by a Wellcome Trust Senior Research Fellowship 202788/Z/16/Z and Medical Research Council (MRC) grants MR/K02213X/1 and MR/L009013/1. Brain samples were provided by the Oxford Brain Bank (BBN004.29852). The Wellcome Centre for Integrative Neuroimaging is supported by core funding from the Wellcome Trust (203139/Z/16/Z). We acknowledge the Oxford Brain Bank, supported by the Medical Research Council (MRC), Brains for Dementia Research (BDR) (Alzheimer Society and Alzheimer Research UK), and the NIHR Oxford Biomedical Research Centre. The views expressed are those of the authors and not necessarily those of the NHS, the NIHR or the Department of Health.

