## Supplementary Material for "Use of multi-flip angle measurements to account for transmit inhomogeneity and non-Gaussian diffusion in DW-SSFP"

### Supporting Figures

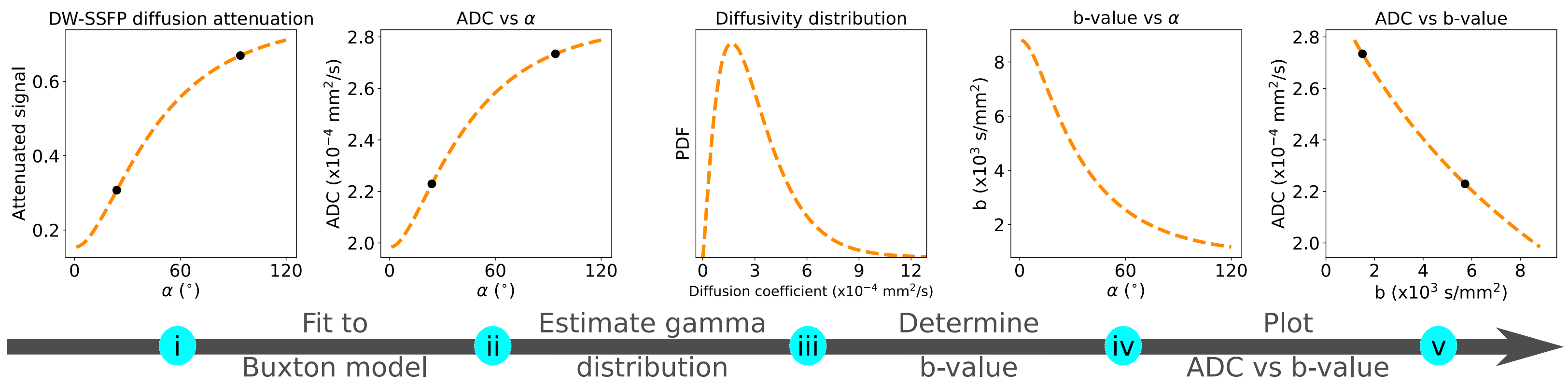


Supporting Figure S1: Outline of the processing pipeline to generate ADC estimates at a single $b_{eff}$. Experimental DW-SSFP data acquired at two flip angles (i - dots) is converted into ADC estimates (ii - dots) using the Buxton signal model (Buxton, 1993). To explain the variation in ADC with flip angle, a DW-SSFP signal model incorporating a gamma distribution of diffusivities (Tendler et al., 2019) is fit to the ADC estimates, to determine a diffusivity distribution (iii) which is able to explain the variation in ADC (ii – dashed line). A DW-SE signal can be subsequently simulated assuming the same diffusivity distribution (iii). From this, we can determine which DW-SE b-value gives rise to the same ADC estimate per DW-SSFP flip angle (iv). The ADC estimates with DW-SSFP can be subsequently plotted vs an DW-SE b-value (v), here defined as  the DW-SSFP effective b-value, $b_{eff}$. By interpolating (or extrapolating) along this curve, we can define an ADC estimate at any $b_{eff}$. By repeating this process in every voxel, we can generate an ADC map at a single $b_{eff}$.

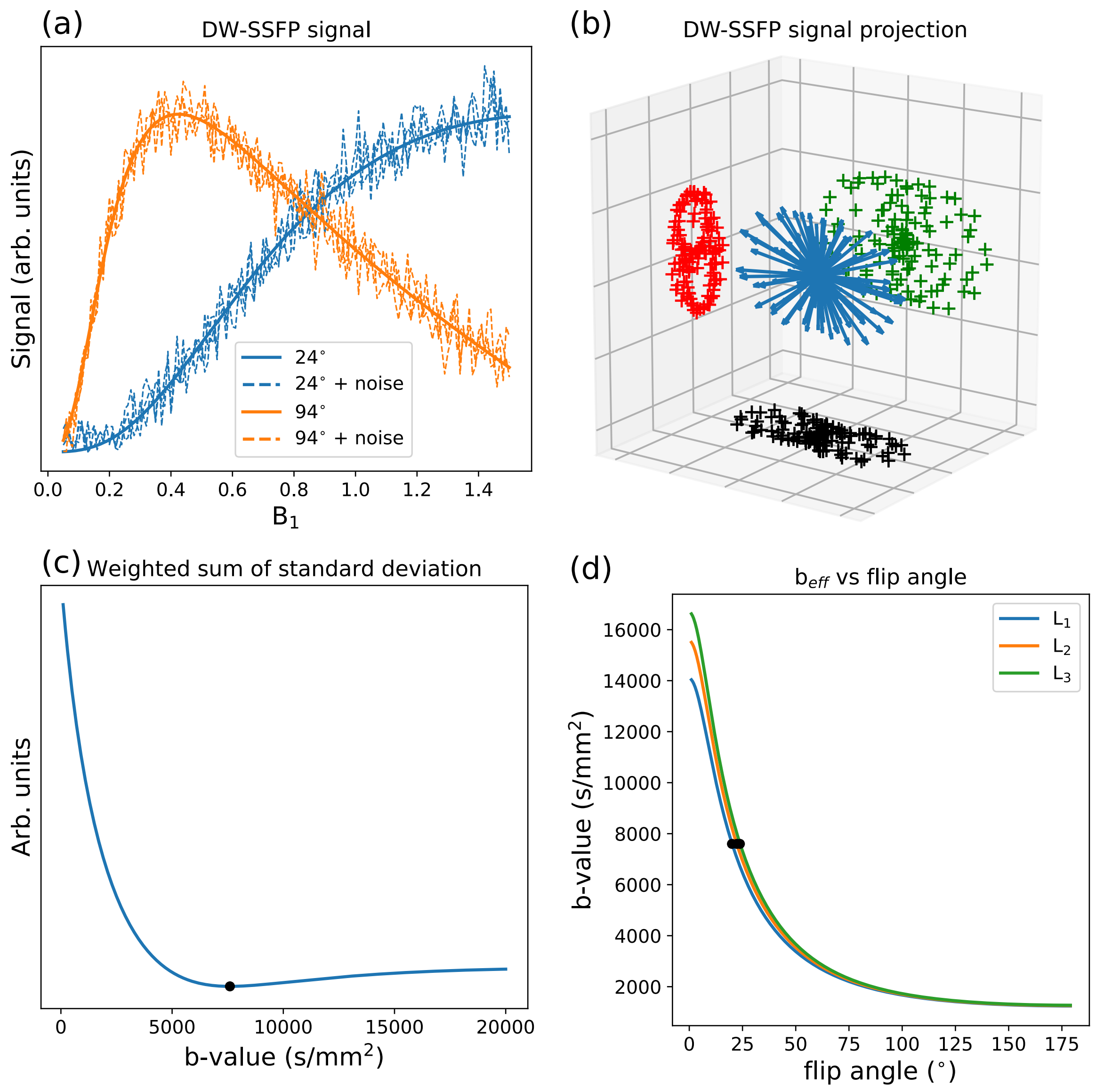


Supporting Figure S2: (a) Example DW-SSFP signal curve simulated at nominal flip angles 24º and 94º. The dashed lines display three different signal curves at each nominal flip angle with added Rician noise. (b) Example distribution of the DW-SSFP signal simulated over the 120 diffusion directions. To highlight the anisotropy of the distribution, the baseline signal common to all 120 diffusion directions was subtracted. (c) Results of the optimization to determine an SNR-optimal $b_{eff}$, determined as $b_{eff} = 7600$ s/mm$^2$ (black dot). (d) Simulating the $b_{eff}$ over a range of flip angles, our SNR-optimal $b_{eff}$ corresponds to a flip angle of $\sim 20^o - 24^o$ over $L_{1,2,3}$.

#### Determination of an SNR-optimal $b_{eff}$

*Theory*

The signal and diffusion-attenuation in DW-SSFP has a strong dependence on flip angle (Figs. 1b and c - main text). This leads to flip angle dependent ADC estimates in systems with non-Gaussian diffusion (Fig. 3 - main text) (Tendler et al., 2019). In our work we have acquired DW-SSFP datasets at two nominal flip-angles to address the issue of variable SNR due to $B_1$ inhomogeneity. However, the resulting eigenvalue maps reveal diffusivity estimates that depend on flip angle (Fig. 8 - main text), preventing a simple averaging.

By modelling the influence of flip angle on ADC, we can define our diffusivity estimates in terms of an effective b-value, $b_{eff}$. Diffusivity maps can be subsequently defined at a single $b_{eff}$ across the entire sample (see Theory – "*A DW-SSFP effective b-value*" in the main text). Importantly, the SNR of the resulting diffusivity maps will depend on the choice of $b_{eff}$. For example, given a set of sample properties and DW-SSFP sequence parameters, different flip angles lead to different $b_{eff}$ estimates (Fig. 3d - main text). Choosing a $b_{eff}$ that corresponds with a flip angle that yields high SNR DW-SSFP data (Fig. 1b - main text) will lead to SNR-optimal diffusivity estimates.

To determine an SNR-optimal $b_{eff}$, we investigated how different $b_{eff}$ estimates affected the SNR of resulting diffusivity maps. This was achieved by simulating the DW-SSFP signal with added Rician noise. Simulations were designed to closely followed the sample properties and acquisition protocol used in our experimental analysis (see Methods – "*MRI Data acquisition protocol*" in the main text).

*Method*

To determine an SNR optimal $b_{eff}$, we performed simulations with synthetic DW-SSFP datasets generated at $24^{o}$ and $94^{o}$ over a range of $B_1$ values with added Rician noise (Fig. S2a). The synthetic datasets were simulated using the Buxton model of DW-SSFP (Buxton, 1993) assuming a diffusion tensor model and a non-Gaussian diffusion estimator defined by a gamma distribution of diffusivities. The synthetic datasets were designed to closely match the properties of the postmortem brain samples and experimental acquisition (see Methods – "*MRI Data acquisition protocol*" in the main text) as follows:

- At each flip angle, DW-SSFP datasets over 120 diffusion directions ($q = 300\ \mathrm{cm}^{-1}$) and six non-diffusion weighted datasets were simulated. Diffusion directions were determined using the GPS tool in FSL (Jenkinson et al., 2012; Jones et al., 1999).
- To match our experimental protocol, $\mathrm{TR} = 28\ \mathrm{ms}$.
- $\mathrm{T}_1 = 574\ \mathrm{ms}$ and $\mathrm{T}_2 = 30.7\ \mathrm{ms}$, the mean values over white matter of all five post-mortem brains.
- The datasets were simulated under a non-Gaussian diffusion estimator (gamma distribution of diffusivities), setting $D_{m1} = 2.988 \cdot 10^{-4}\ \mathrm{mm}^2/\mathrm{s}$, $D_{m2} = 2.254 \cdot 10^{-4}\ \mathrm{mm}^2/\mathrm{s}$, $D_{m3} = 1.786 \cdot 10^{-4}\ \mathrm{mm}^2/\mathrm{s}$, $D_{s1} = 4.044 \cdot 10^{-4}\ \mathrm{mm}^2/\mathrm{s}$, $D_{s2} = 3.158 \cdot 10^{-4}\ \mathrm{mm}^2/\mathrm{s}$ and $D_{s3} = 2.538 \cdot 10^{-4}\ \mathrm{mm}^2/\mathrm{s}$ the mean values of $D_m$ and $D_s$ along $L_{1,2,3}$ over white matter estimated from the five post-mortem brains. The orientations of the eigenvectors $\vec{V}_1$, $\vec{V}_2$ and $\vec{V}_3$ were kept constant for all simulated signals.
- The datasets were simulated over the range $B_1 = 0.05$ to $B_1 = 1.50$, in steps of 0.01. For each $B_1$ value, 1000 noise repeats were simulated, with the noise level estimated from the repeats of the experimental non-diffusion weighted DW-SSFP data (Fig. S2a).

An example simulated DW-SSFP signal distribution over all 120 directions is displayed in Fig. S2b.

The simulated data were subsequently processed in an identical manner to the two-flip angle experimental data as follows:

- Unique $L_{1,2,3}$ estimates were generated at 24° and 94° for each $B_1$ value and noise repeat (see Methods – "*Data processing*" in the main text)
- $D_{m1,2,3}$ and $D_{s1,2,3}$ were subsequently determined from the $L_{1,2,3}$ estimates at 24° and 94° (see Methods – "*Combination of eigenvalue estimates at two-flip angles to a single effective b-value*" in the main text).
- From the values of $D_{m1,2,3}$ and $D_{s1,2,3}$, $L_{1,2,3}$, estimates were generated for a single $\mathrm{b_{eff}}$, over the range $\mathrm{b_{eff}} = 100\ \mathrm{s/mm}^2$ to $\mathrm{b_{eff}} = 20000\ \mathrm{s/mm}^2$ in steps of $100\ \mathrm{s/mm}^2$.

From the $L_{1,2,3}$ estimates simulated at each $\mathrm{b_{eff}}$, the standard deviation was determined over the 1000 noise repeats at each $B_1$ value. The standard deviations estimated at each

$B_1$ value were subsequently weighted by the $B_1$ histogram of the experimental data (Fig. 5 – main text) and summed. The $b_{eff}$ with the minimum summed standard deviation was defined as the SNR-optimal $b_{eff}$.

*Results*

Figure S2c reveals the distribution of the weighted sum of the standard deviation with $b_{eff}$, determining the SNR-optimal $b_{eff} = 7600\ s/mm^2$ (Fig. S2c – black dot). As our simulations have been performed for a set of fixed parameters, each flip angle will correspond to a unique $b_{eff}$ along $L_{1,2,3}$. Plotting the relationship between $b_{eff}$ and flip angle (Fig. S2d), we can determine that the SNR-optimal $b_{eff}$ corresponds to a flip angle of $20^o - 24^o$ (Fig. S2d – black dots). Considering the $B_1$ profile of all 5 postmortem brain samples, approximately 82%-93% of voxels are being interpolated to $b_{eff} = 7600\ s/mm^2$, with the remaining voxels extrapolated.
